# Correlations among hydrogen bond fluctuations in the apo state are enough to reveal allosteric networks in proteins

**DOI:** 10.1101/2025.04.23.650155

**Authors:** Subhajit Sarkar, Saikat Dhibar, Biman Jana

## Abstract

Allostery is essential for regulating biochemical pathways, where ligand binding at one site influences enzymatic activity at a distant functional site. Identifying allosteric sites and mapping signal transduction pathways in biomolecules remain a significant challenge. Existing experimental or computational methods, in general, can identify a subset of total allosteric network. Here, we consider that in addition to providing structural stability and flexibility of proteins, hydrogen bonds inside the protein may act as conduits for long-range communication. To this end, we develop a computational framework that predicts allosteric sites and total allosteric network by analyzing correlations among hydrogen bond fluctuation from equilibrium molecular dynamics trajectory of apo state of proteins. We demonstrated that this method can accurately captures experimentally verified allosteric sites and suggest allosteric signal transduction pathways across three different proteins. Furthermore, since our predictions are derived solely from the simulation trajectory of the apo state, these findings reinforce the idea that the signature of allostery is inherently encoded in the apo state of the protein. This approach offers a useful strategy to decode allosteric network and pockets, with broad implications for drug discovery and the targeted modulation of allostery in proteins.

**Significance:** Allostery, distal ligand-induced functional modulation of biomolecules, is central to biological regulation, yet its intricate mechanisms remain elusive. Precise identification of allosteric sites and pathways is essential for targeted drug development, minimizing off-target effect. Here, we introduce a novel computational approach, Hydrogen Bond Allosteric Map (HBAlloMap), which leverages the dynamic fluctuations of hydrogen bond networks derived from microsecond molecular dynamics simulations to comprehensively map allosteric modules within proteins. By analyzing the fluctuation of correlated hydrogen bonds, our method effectively reveals key allosteric hotspots and signal transduction pathways in PDZ3, PDZ2, and Pin1. By providing a robust and efficient tool for deciphering allosteric mechanisms, this method has the potential to accelerate drug discovery and deepen our understanding of allostery.

---

Allostery (1,2) is a fundamental regulatory mechanism where the binding of a ligand at one site of a biomolecule influences the activity of a distant functional site. Often referred as “second secret of life,” it plays a crucial role in enzyme regulation (3), receptor signaling (4), and biochemical signal transduction(5) and has far-reaching implications in pharmacological research and drug discovery(6, 7). Despite its central role in biology, understanding how local perturbations such as ligand binding propagate through the protein structure to modulate the function of remote sites still remains a mystery (8–11).

Classical models of allosteric regulation, such as the Monod-Wyman-Changeux (MWC)(12) and Koshland-Némethy-Filmer (KNF)(13) models, propose that ligand binding triggers significant structural changes in a protein. These conformational shifts drive functional modulation by stabilizing distinct states. This form of conformational allostery is well-established in systems like hemoglobin, where cooperative binding relies on large-scale structural rearrangements.

These models provided a foundation for understanding allosteric transitions as discrete conformational changes, directly linking structural shifts to functional regulation. However, recent evidence challenges the traditional structure-centric view of allostery, showing that allosteric regulation can occur without significant conformational changes. This has led to the concept of dynamic allostery (14–17), where functional modulation arises from shifts in protein dynamics rather than large structural transitions. In this mechanism, ligand binding perturbs the protein’s intrinsic flexibility, altering the dynamic equilibrium of side chains and backbone motions (2). These changes affect conformational entropy, redistributing the populations of pre-existing states to drive allosteric communication. In systems such as PDZ domain(14), Pin1(18) and met repressor(17), presence of dynamic allostery have been reported.

Identifying allosteric sites in proteins is a challenging task. Both theoretical and experimental approaches have been developed to detect these sites with greater accuracy. Precise identification of allosteric sites is crucial in drug development, as targeting these regions can significantly influence therapeutic strategies. Unlike orthosteric drugs, which bind to the active site and often lead to serious side effects due to conserved structural features across protein families, allosteric drugs target less conserved regions. This results in higher specificity, reduced off-target effects, and finer control over protein function (19). These advantages make allosteric drugs a preferred choice over their orthosteric counterparts in many therapeutic applications. To unravel the allosteric site and allosteric signal transduction pathways, structural and energetic fluctuation correlation analyses have previously been extensively employed (20–22). Computational approaches have played a pivotal role in deciphering the allosteric mechanisms. Methods such as statistical coupling analysis (SCA) (23) and direct coupling analysis (DCA)(24) identify co-evolving residues that contribute to allosteric signaling networks. Advanced techniques, including deep coupling scan (25) and rigid-residue scan (RRS)(26) provide insights into the residue-specific contributions to the propagation of allosteric signals. Anisotropic thermal diffusion (ATD) (27) method models the directional transfer of thermal fluctuations, offering a detailed understanding of how energy flows through the protein structure. Similarly, interaction correlation method(20) identified energetically correlated protein regions shedding light on the energetic and structural interdependencies that facilitate allosteric communication. Modern machine learning (28, 29) and deep learning methods (30) have also been used to decrypt allosteric pockets and to decipher allosteric signal transduction pathways in proteins. Among experimental techniques, nuclear magnetic resonance (NMR) stands out for its ability to reveal both the structural and dynamic properties of biomolecules. NMR relaxation experiments provide valuable insights into protein dynamics, enabling the detection of allosteric processes in diverse biomolecular systems (31, 32). When combined with binding assays and site-directed mutagenesis, NMR can identify key residues involved in the transmission of allosteric signals, offering a detailed understanding of allosteric communication pathways. Additionally, techniques such as double mutant cycle(33) have been employed to map energetic networks within proteins, enabling the characterization of allosteric mechanisms across different systems. These experimental approaches allow for the comparative analysis of homologous proteins and contribute significantly to elucidating the fundamental principles underlying allostery. While each of these methods identifies a part of the whole allosteric module of the protein, it comes with some noise. Therefore, combination of these existing methods may illuminate the whole allosteric module of the protein, but such data will also have significant noise in it. Thus, there is a real need to develop a method which can provide the most parts of the whole module at once with minimal noise.

In this article, we present Hydrogen Bond Allosteric Map (HBAlloMap), a method that can identify the whole allosteric module of a protein. Our method uses the fluctuation correlation of all possible hydrogen bonds in a protein from microsecond long molecular dynamics trajectory at the apo state. We have demonstrated that our method can identify the allosteric hotspots and provide suggestions on the possible allosteric signal transduction pathways in three model systems, PDZ3, PDZ2 and Pin1. Therefore, this new method can be used for other systems to identify the allosteric sites/pockets which can advance the drug design research further.

## Results and Discussion

Method for identifying allosteric hotspots: Hydrogen Bond Allosteric Map (HBAlloMap)

Hydrogen bonding interactions are known to provide a significant part of energetics of a protein molecule along with other contributions (43). Hydrogen bonds are weak, non-covalent interactions formed between a hydrogen atom covalently bonded to an electronegative atom (typically nitrogen, oxygen, or fluorine) and another nearby electronegative atom (44). Despite being weaker than covalent bonds, hydrogen bonds are highly directional and relatively easy to break, making them essential for maintaining protein stability, facilitating conformational changes, and supporting functional adaptability. Their unique balance of strength and flexibility allows them to stabilize structural intermediates while remaining dynamic enough to break and reform during conformational transitions. Moreover, hydrogen bond networks may span considerable distances within proteins, enabling the propagation of allosteric signals across different domains. In this study, we have developed a hydrogen bond-based method capable of accurately predicting experimentally verified allosteric sites and delineating allosteric signal transduction pathways in systems which show dynamic allostery.

We have performed two microsecond molecular dynamics simulation for each of the apo proteins and track each of the intra-protein hydrogen bond present over the simulation trajectory. Next, we have constructed hydrogen bond time series matrix consisting of all possible hydrogen bonds present in the molecular dynamics simulation data. The donor-acceptor distance cutoff was 3.5 Å and donor-hydrogen-acceptor angle cutoff was set at 30 degree for hydrogen bond calculation. If a particular hydrogen bond is present in a particular simulation timeframe it is considered as 1, otherwise 0 is assigned to that timeframe. Therefore, a configuration of protein is described as a vector having components either 0 or 1. Covariance analysis of the hydrogen bond time series matrix was performed. Then diagonalization of the matrix was performed followed by eigen-decomposition. Highest eigenvalue components (mainly the principal component 1) were considered, the absolute values of the respective components of the eigenvectors of these principal components were taken in a descending order. The residues of hydrogen bonds having larger absolute values were decorated in the protein structure. A schematic workflow of Hydrogen Bond Allosteric Map (HBAlloMap) is presented in Figure 1.

**Figure 1:**
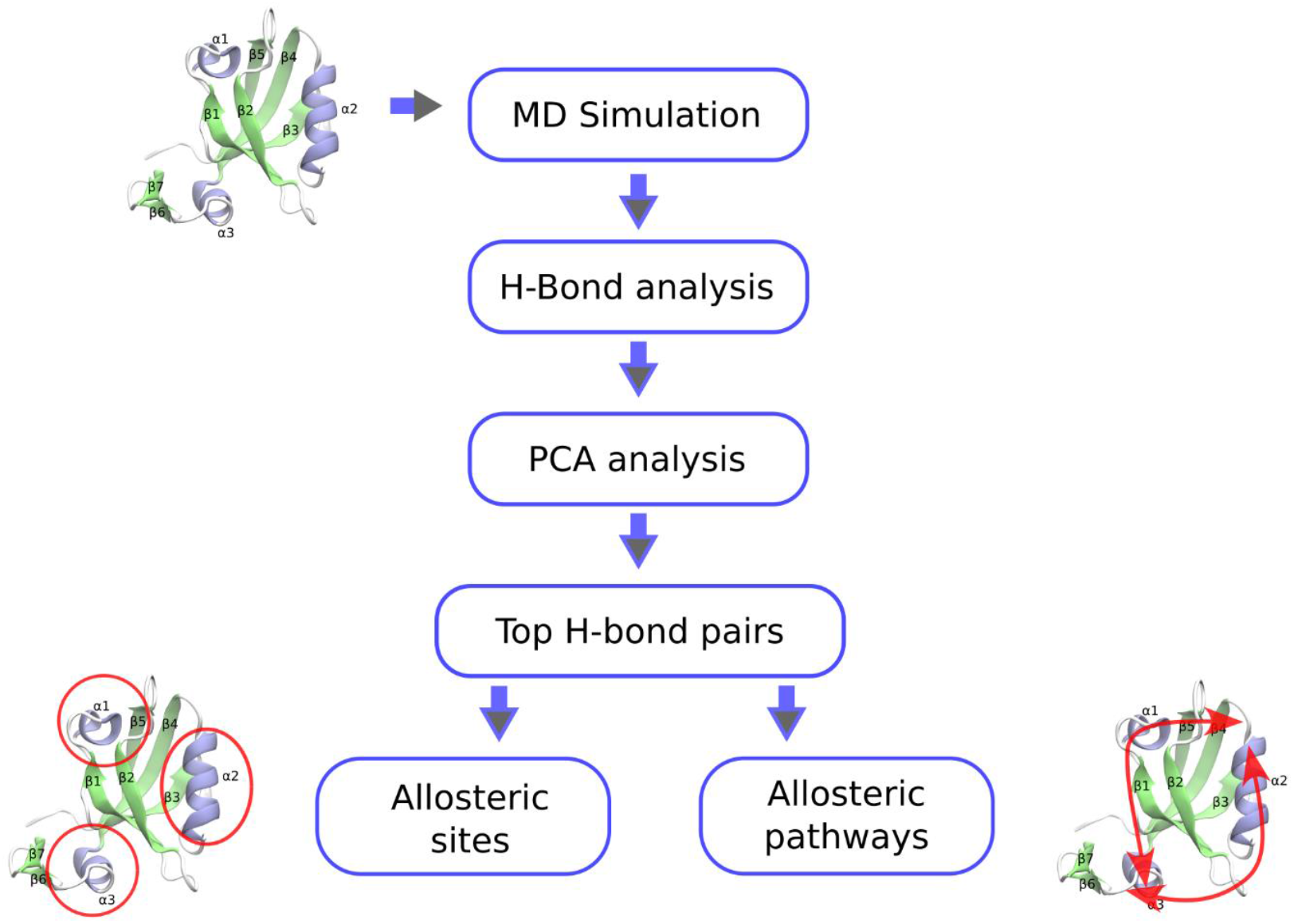
Schematic representation of HBAlloMap method. The diagram illustrates the key computational steps involved in detecting functionally important regions and mapping long-range allosteric communication.

### HBAlloMap detects residues important for allosteric communication in PDZ3 domain

The PDZ3 domain is an important protein–protein interaction module that helps regulate signaling pathways and scaffolding functions (45). PDZ3 domain has been widely studied as a model system for single-domain dynamic allostery as it has a clearly identifiable binding and allosteric site. First discovered in synapses of the mammalian nervous system (46), it is one of the most common protein domains in eukaryotes. PDZ3 has a conserved structural fold made up of six β-strands and three α-helices. Its binding pocket, formed by the βB strand and αB helix, allows ligand peptides to bind in an antiparallel β-strand conformation relative to βB (34) (Figure 2a). This domain mainly binds to the C-terminal tails of target proteins, recognizing short sequences of four to five residues, with specificity determined by sequence motifs.

**Figure 2.**
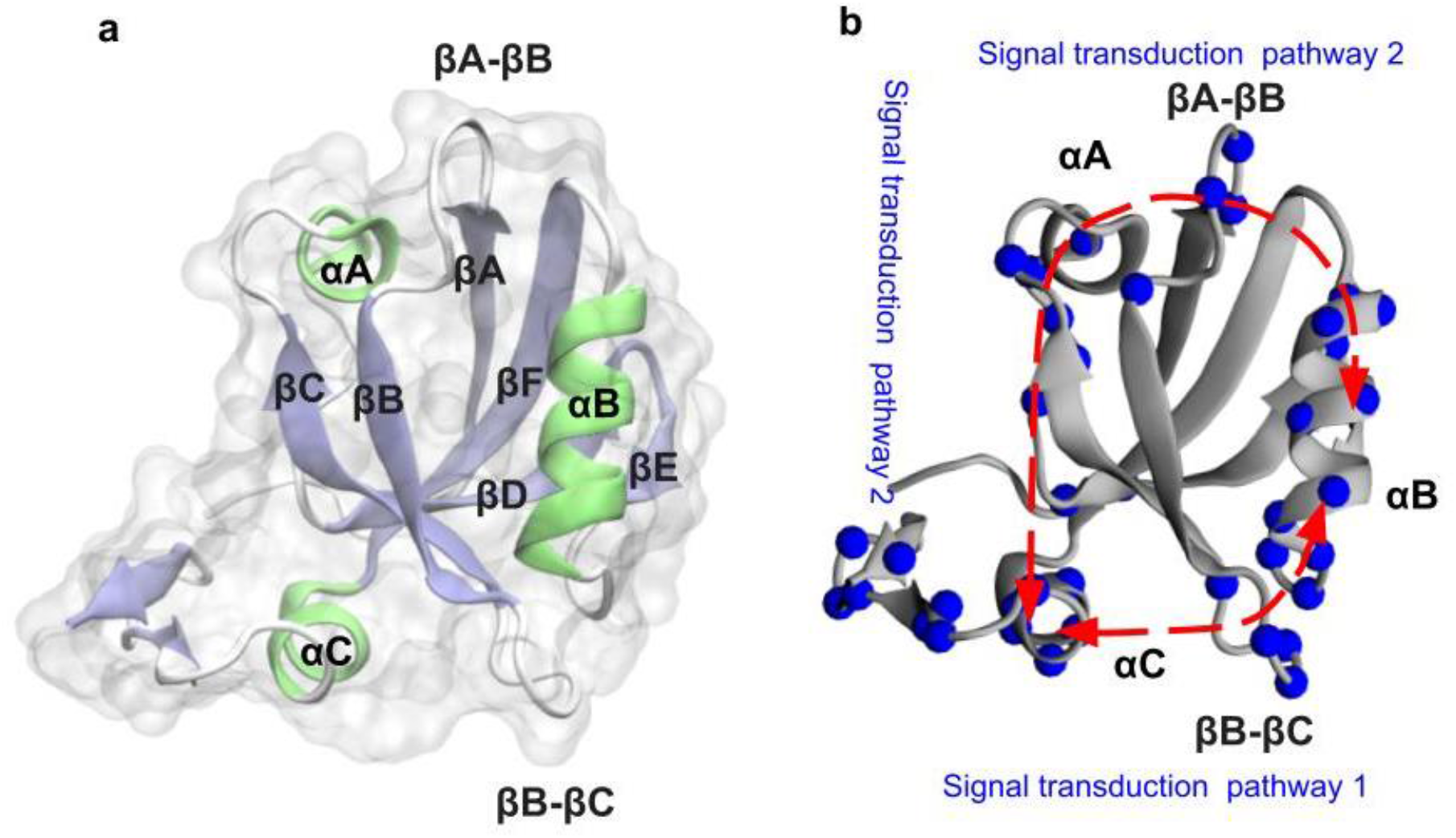
(a) Crystal structure of the apo state PDZ3 domain from PSD-95 (PDB ID: 1BFE), with key secondary structure elements highlighted. α-helices are shown in lime, while β-strands are depicted in iceblue, illustrating the overall fold of the domain. A semi-transparent surface representation provides structural context, emphasizing the spatial arrangement of these elements.(b) Cα atoms of residues of the first 39 unique residues (35 percent of total number of protein residues) obtained from first principal component of hydrogen bonds are decorated on the apo state crystal structure of PDZ3 (PDB ID 1BFE). As it can be seen, the predicted residues successfully captures the experimentally verified allosteric site (αC helix), and two different pathways from the distant allosteric site to the binding site emerges. In figure 2b these two pathways are marked as signal transduction pathway1 and pathway2.(residue lists with respect to each pathway is given in Note S1)

We have performed two microsecond simulation of apo state PDZ3 (PDB 1BFE). From apo state simulation we have constructed hydrogen bond time series data by tracking each of the hydrogen bond over the simulation trajectory. We have done PCA (Principal Component Analysis) on that hydrogen bond time series data. Following our protocol, we highlight the first 39 unique residues (35 percent of total number of residues) obtained from first principal component of hydrogen bonds on the structure as depicted in Figure 2b. From Figure 2b it can be seen that this method captures the αC helix (GLU 401,ARG 399,SER 398,TYR 397,GLU 396,GLU 395), αB helix (SER 371,HIS 372,GLU 373,ALA 376,ILE 377, LYS 380, ASN 381, ALA 382), βA-βB loop(THR 321,GLY 319,ARG 318), βB-βC loop (GLY 335,GLU 331, ASP 332, GLY 333), and αA helix and nearby loop (ARG 354, LEU 353, GLU 352, GLY 351, SER 350,ALA 343) (residue lists are attached in the Table S1). Next, we will discuss about the previous experimental/theoretical evidences for these regions in terms of allosteric communication in PDZ3 domain.

Previous experimental and theoretical studies have identified the αC helix as a crucial regulator of allostery in the PDZ3 domain. Petit et al. investigated its role by comparing the wild-type PSD-95 PDZ3 to a truncated version lacking the αC helix. Using NMR, they found that removing residues of αC helix (396–402) did not significantly alter the overall fold of PDZ3. However, it caused a 21-fold reduction in binding affinity, highlighting the αC helix’s role in stabilizing ligand interactions (14). Zhang et al. employed NMR experiments to investigate phosphorylated PSD-95 PDZ3 domain. Their findings supported the notion of αC helix having a crucial role in peptide binding affinity of PSD-95 PDZ3 (47). Bozovic et al. engineered a photocontrollable PDZ3 domain by incorporating an azobenzene-based photoswitch into the αC-helix. Their findings showed that light-induced structural perturbations in the αC-helix led to a substantial shift in binding affinity, reaching up to ∼120-fold (48). Computational studies have also supported the role of the αC helix in controlling the ligand binding affinity of PDZ3 domain. Rigid-body molecular dynamics simulations done by Kalescky et al. identified residues of the αC helix as important for PDZ3 allostery(26). Kumawat et al. used classical MD simulations to identify the allosteric network by decomposing the average pairwise interaction energies among all residues of the PDZ3 domain and analyzing their perturbations upon ligand binding. This analysis revealed a distinct network that connects the ligand-binding site to the distal allosteric site, with the αC helix serving as a critical conduit for allosteric signal propagation(49).

Normal mode analysis (NMA) of PSD-95 PDZ3 revealed that a shift in the αB helix expands the binding pocket, facilitating ligand binding (50). Conservation-mutation correlation analysis (CMCA) identified the αB helix as part of a correlated network involved in allosteric communication (51). Perturbation response scanning (PRS) further supported its role by showing that applied forces on the protein caused structural displacements in the αB helix, indicating its involvement in allosteric propagation (52). A single-mutation scan combined with statistical coupling analysis (SCA) linked the αB helix to ligand binding, reinforcing its functional importance (53). Molecular dynamics (MD) simulations by Morra et al. demonstrated significant energetic modulations in the αB helix upon ligand binding (21). Similarly, Kalescky et al. used non-equilibrium rigid-body MD simulations and identified the αB helix as a key element in global allosteric signal transmission (26). These findings provide strong theoretical and experimental evidence supporting the αB helix as a critical allosteric hotspot in PDZ3.

In addition to αC helix, αβ helix and adjacent lower loop, our method also identifies βB – βC (GLY 335,GLU 331, ASP 332, GLY 333) and βA – βB (THR 321, GLY 319,ARG 318) loop as potential allosteric hotspot (residue lists are attached in Table S1). In our previous work, we have shown that these two distant loops show correlated dynamics in the wild-type PDZ3 domain, which vanished upon αC helix truncation/mutation. These correlated dynamics are central to allosteric motion shown by PDZ3 domain(54).

Our method also captures some residues adjacent to αA helix (ARG 354,LEU 353,GLU 352, GLY 351, SER 350, ALA 343) (see Table S1). Computational techniques like Anisotropic thermal diffusion (27), Rotamerically induced perturbation(55), Conservative –mutation correlation analysis(51), Perturbation response scanning(52) have pointed to the role of αA helix in PDZ3 allostery.

Finally, please note that while different existing experimental and computational methods have identified different structural regions of the PDZ3 domains allosteric communication, HBAlloMap actually reveal all those regions at once.

### Suggestion of allosteric signal transduction pathway of PDZ3 domain

From the HBAlloMap results we have seen that important allosteric site residues can be captured. Based on our result, we have also tried to suggest the allosteric signal transduction pathways. As can be seen from figure 2b, two different pathways from the distant allosteric site (αC helix) to the binding site emerges. In figure 2b these two pathways are marked as signal transduction pathway 1 and 2. Pathway1 starts from the αC helix (experimentally verified allosteric site) and consists of βB-βC loop, αB helix to the binding site, which is in accordance with our previous observations(54). Pathway 2 starts from the αC helix and consists of αA helix and nearby loop and βA-βB loop ending in canonical binding site (residue mapping with respect to each pathway is given in Note S1). Similar kind of pathway like pathway 2 was predicted by Kong et.al (20) using Interaction correlation analysis method in a different PDZ domain (PDZ2 domain from human phosphatase HPTP1E). Please note that these pathways/ residues are found from the apo state simulation of PDZ3 which indicates that probably allosteric signal network is inscribed even in the apo state protein dynamics.

### HBAlloMap detects experimentally verified allosteric sites and suggests allosteric signal transduction pathways in PDZ2 domain

Next, we have performed HBAlloMap method on two microsecond simulation trajectory of apo state of PDZ2 domain (PDB ID 3LNX). The PDZ2 domain is essential for various cellular functions, including cell-cell communication, polarization, and protein regulation (56). It plays a key role in forming and stabilizing protein complexes, linking extracellular signals from transmembrane receptors to intracellular responses. This connection is vital for maintaining cellular organization and function. Following our protocol, we have extracted the first principal component from the hydrogen bond time series data. Next, we highlight 33 top unique residues (35 percent of total number of protein residues) obtained from first principal component on the structure of PDZ2 as depicted in figure 3b, our method clearly identifies the αB helix (GLN 83,ASN 80,ARG 79, GLU 76,VAL 75,GLN 73,THR 70), αA helix and nearby loop (ARG 51,ASP 49,SER 48,GLU 47), βA –βB loop (SER 17,ASN 16,ASP 15), βB-βC loop (ASN 27,THR 28,VAL 30,ARG 31,HIS 32) as allosterically important sites in PDZ2 (residue lists are attached in the Table S2). Next we discuss about the available experimental/computational evidences on these structural region of PDZ2 in terms of allosteric communication.

**Figure 3:**
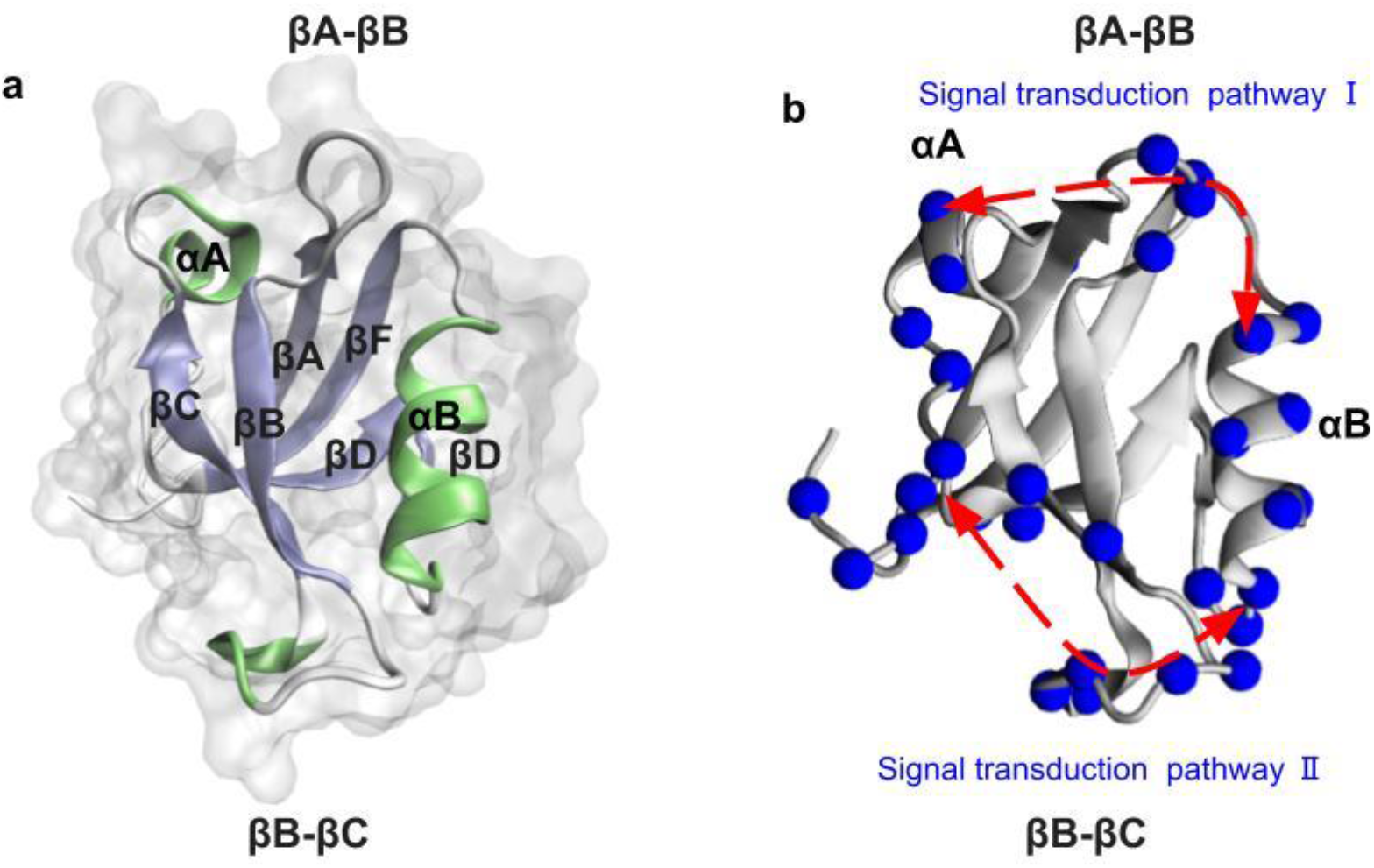
(a) Crystal structure of the PDZ2 domain from human PTP1E(PDB ID: 3LNX), with key secondary structure elements highlighted. α-helices are shown in lime, while β-strands are depicted in iceblue, illustrating the overall fold of the domain. A semi-transparent surface representation provides structural context, emphasizing the spatial arrangement of these elements.(b)Cα atoms of residues of the first 33 unique residues (35 percent of total number of residues) obtained from first principal component of hydrogen bonds are decorated on the apo state crystal structure of PDZ2(PDB ID: 3LNX). As it can be seen, the predicted residues successfully captures the experimentally verified allosteric site (αA helix), and two different allosteric signal transduction pathways emerge. In figure 3b these two different allosteric signal transduction pathways are designated as I and II.

**Figure 4:**
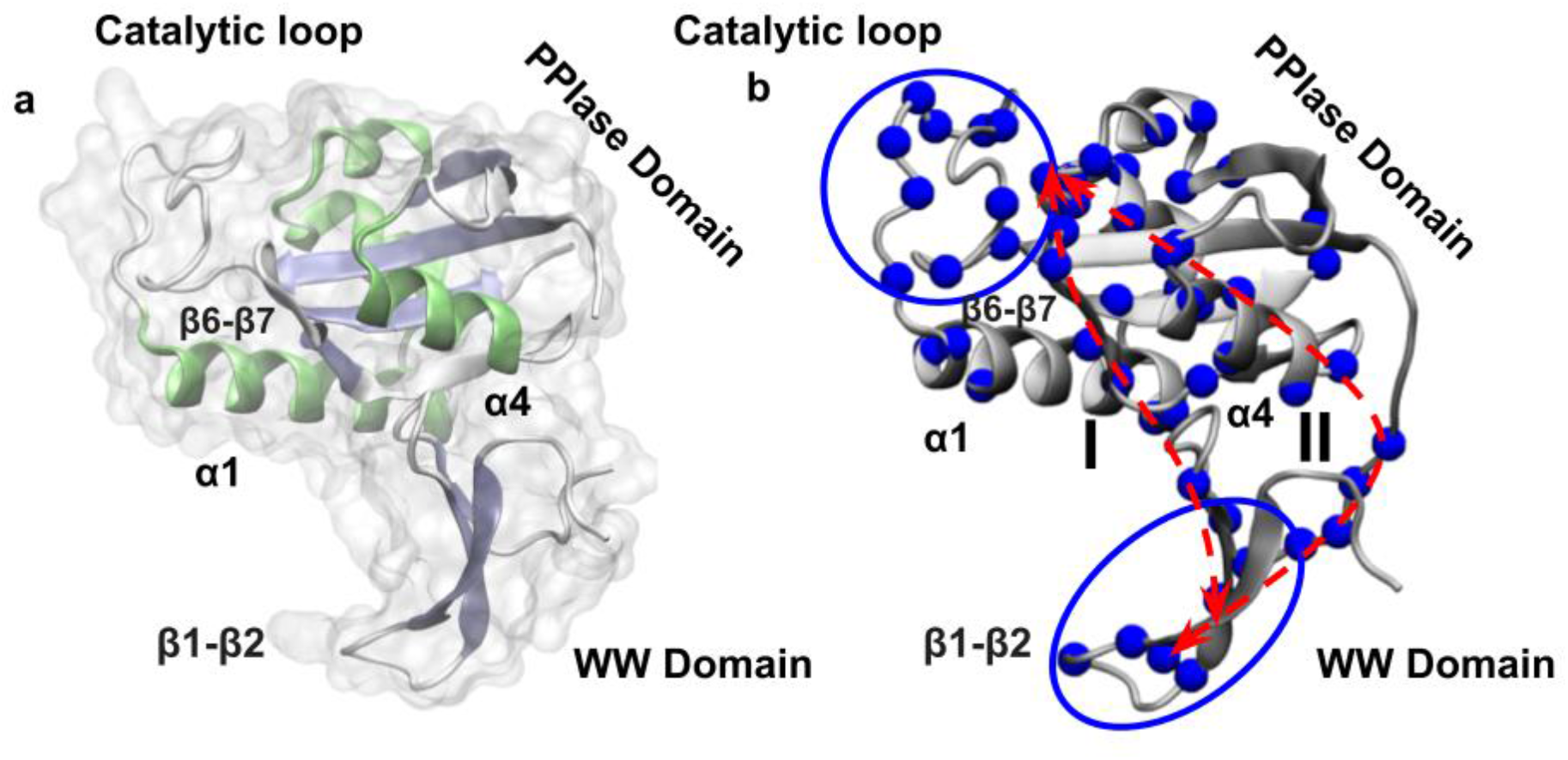
(a) Crystal structure of Pin1, a multidomain protein consisting of an N-terminal WW domain and a C-terminal PPIase domain. Key secondary structural elements are highlighted, with α-helices in lime and β-strands in iceblue, emphasizing their spatial arrangement. Important regions involved in allosteric regulation, including the catalytic loop, β6-β7 strands, α1, and α4 helices, are labeled. A semi-transparent surface representation provides additional structural context.(b) Cα atoms of residues of first 55 unique residues (35 percent of total number of protein residues) obtained from first principal component of hydrogen bonds are decorated on the apo state crystal structure of Pin1 (PDB ID 3TBD). As it can be seen, predicted residues successfully captures the experimentally verified allosteric site (WW domain)and binding site, two different pathways from the distant allosteric site to the binding site emerges. In figure 4b these two pathways are marked as pathway I (WW domain -interdomain interface-PPIase domain core (α4 helix and β6-β7 loop region) –catalytic loop region) and pathway II (WW domain - PPIase peripheral α1 helix, and the α1 helix-core interface to the catalytic loop). For detailed residuewise mapping of signaling pathway see Note S3.

Previous studies have identified dynamic allostery in the αB helix and its adjacent lower loop using both experimental and computational methods. Experimental techniques such as NMR chemical shift analysis by Fuentes et al. (31), point mutations with NMR by Fuentes et al.(57), and equilibrium MD simulations with NMR restraints by Dhulesia et al.(22) have all recognized the αB region as an allosteric hotspot.

Computational studies further support these findings. Equilibrium MD simulations by Kong et al.(20), and Morra et al.(21) have identified the αB helix and its adjacent lower loop in PTP-BL PDZ2 as key allosteric regions. In addition to traditional simulations, a variety of other computational approaches including perturbation response scanning (PRS) (52) and elastic network models (ENM) (58), also highlight the αB helix and adjacent loop as critical elements in PDZ2 allosteric regulation.

NMR experiments have shown that ligand binding triggers allosteric changes in the αA helix of the PDZ2 domain. Walma et al. (59) detected chemical shifts in the αA helix of PTP-BL PDZ2 upon binding to the human Fas receptor and RIL C-terminal peptides. Fuentes et al. observed dynamic changes in this region when bound to the RA-GEF2 peptide (31). Computational studies have successfully reproduced these findings. Techniques such as Monte Carlo sampling (60), perturbation response scanning (PRS) (52), and anisotropic thermal diffusion (ATD) (27) have all confirmed the αA helix as a key site of allosteric regulation in PTP-BL PDZ2.

In addition to αB and αA helix region our method also captures βA-βB (SER 17,ASN 16,ASP 15) and βB –βC (ASN 27,THR 28,VAL 30,ARG 31,HIS 32) loop region as potential allosteric sites (residue list for these hotspots are attached in Table S2). NMR relaxation experiments done by Fuentes et. al (57) and Dhulesia et. al (22) also finds some of the residues of these two loop regions as important potential allosteric sites.

Here also, while different experimental and computational techniques find one or the other region responsible for allostery in PDZ2, our method seems to find all of them at once.

### Suggestion of allosteric signal transduction pathways of PDZ2 domain from HBAlloMap

From the results of HBAlloMap, we have seen that important allosteric site residues can be captured for PDZ2 domain. We have tried to suggest the possible allosteric signal transduction pathways based on our result. Pathway I connects the αB helix to the distal αA helix via βA –βB loop and pathway II connects the αB helix to the distal N terminal region (figure 3b) via βB –βC loop (detailed residuewise mapping of signaling pathway is attached in Note S2). Similar kind of allosteric signal transduction pathways were suggested by Kong et.al. using Interaction correlation method(20).

### Allosteric residues and pathway prediction in Pin1 domain using HBAlloMap

Pin1 is an important therapeutic target for cancer and Alzheimer’s disease (61–63). It consists of an N-terminal WW domain (residues 1–39) and a C-terminal peptidyl-prolyl isomerase (PPIase) domain (residues 50–163), connected by a flexible linker (residues 40–49)(64). The PPIase domain includes a core region (α4-helix and β4–β7 sheets), α1–α3 helices, and a semi-disordered catalytic loop. While both domains recognize phospho-Ser/Thr-Pro substrate motifs, only the PPIase domain catalyzes peptidyl-prolyl isomerization(65, 66). Interestingly, the isolated PPIase domain has a binding affinity nearly 100 times weaker than the full-length Pin1(65–67). This suggests that the noncatalytic WW domain can allosterically regulate the catalytic activity of the PPIase domain.

We have performed HBAlloMap on Pin1 system and it captures experimentally verified allosteric site WW domain (MET 15, SER 16, SER 18, ARG 21,TYR 23, TRP 34, GLU 35, ARG 36, PRO 37, SER 38), the catalytic loop (ARG 68, SER 72, ARG 74, GLN 75, GLU 76, LYS 77,ILE 78)and functionally important PPIase domain (ARG 142, ALA 140, ALA 137, GLU 135, LYS 132, GLN 131,THR 152, ASP 153, SER 154). First 55 unique residues (35% of total residues) from PC1 captures pathway I which consists of WW domain backside-interdomain interface-PPIase domain core (α4 helix and β6-β7 loop region)–catalytic loop region. It also captures another pathway which consists of WW domain -PPIase peripheral α1, and the α1-core interface to the catalytic loop (pathway II) (detailed residuewise mapping of signaling pathway is attached in Note S3).

### Comparison with previous studies

The two predicted pathways using our method are in accordance with previously predicted pathways by Guo et. al (68) and Xu et. al(30) .Guo et. al use molecular dynamics simulation to propose two different pathways (Pathway I emanates from the WW backside and propagates through the inter-domain interface and the PPIase domain core to the β5-α4 and β6-β7 loops; Pathway II emanates from the WW front pocket and propagates through the bound substrate, the PPIase peripheral α1, and the α1-core interface to the catalytic loop) of allosteric signal transduction in Pin1. Xu et. al used Neural Relational Inference (NRI), based on a graph neural network (GNN), to study and predict allosteric interactions and signal transduction pathways in Pin1. They also predict two different pathways, one consists of WW domain –PPIase core – catalytic loop and other one is WW domain-α1 helix-α2/α3 helix-catalytic loop.

## Conclusions

We have demonstrated that our method can detect important structural region of the proteins which is crucial for allosteric signal transduction. As our method relies only on the correlation of different hydrogen bonds present in the system, it essentially infers that proteins may be relying on the dynamically connected hydrogen bonding network for allosteric signal transduction. These results also suggest that proteins can be considered as a soft material with strongly connected hydrogen bonding network within itself. Our method also provides suggestions on the allosteric signal transduction pathways which can enhance our understanding of information transfer inside biomolecules further. More importantly, our method provides all the structural region involved in the allostery at once which was lacking for other existing methods. Additionally, as we have predicted the allosteric sites and allosteric signal transduction pathways entirely from the simulation trajectory of unbound state, these results profound the idea that signature of allostery is already inscribed into apo state of the proteins. Our method can essentially identify the unknown allosteric structural regions/pockets in important proteins that can enrich the field of allosteric drug design.

## Materials and Methods

### Simulation Details

The crystal structures of PDZ3, PDZ2, and Pin1 were retrieved from the Protein Data Bank (PDB) with accession codes 1BFE (34), 3LNX(32), and 3TDB(35), respectively. All-atom molecular dynamics (MD) simulations of the apo states with explicit solvent were performed using GROMACS 2018.3(36), employing the Amber 99SB-ILDN(37) force field in combination with the TIP4P-Ew (38) water model. To achieve charge neutrality, appropriate amounts of Na+ and Cl− ions were added to each system. The equations of motion were integrated using the leap-frog algorithm with a 2-femtosecond (fs) time step. Initial energy minimization of the structures was carried out using the steepest descent algorithm, followed by equilibration under constant volume (NVT) and constant pressure (NPT) conditions.

Temperature was maintained at 300 K using the velocity-rescaling (39) thermostat with a coupling constant of 0.1 picoseconds (ps), while pressure was controlled at 1 bar using the Parrinello–Rahman barostat (40). Long-range electrostatic interactions were calculated using the particle mesh Ewald (PME)(41) method, and short-range electrostatic and van der Waals interactions were truncated with a cutoff distance of 1.0 nanometer (nm).The Visual Molecular Dynamics package (42) was used to visualize the systems and simulation trajectories.

## Data, Materials, and Software Availability

All study data are included in the article and/or supporting information. Molecular dynamics simulation trajectory files and codes are available upon request.

## Acknowledgements

This research is supported by the Department of Science and Technology SERB grant (Grant ID: C.R.G./2020/000756). The authors extend their sincere gratitude to the central supercomputing facility at the Indian Association for the Cultivation of Science, Kolkata, for providing computational resources. SS is thankful to the Council of Scientific and Industrial Research (CSIR) for awarding a fellowship.

## Supporting Information (SI)

### This pdf file includes

1. Note S1. Predicted allosteric signal transduction pathways for PDZ3
2. Note S2. Predicted allosteric signal transduction pathways for PDZ2
3. Note S3. Predicted allosteric signal transduction pathways for Pin1

**Table S1.** First 39 unique residues (35 percent of total number of protein residues) obtained from first principal component of hydrogen bonds for PDZ3 domain.

| Residue<br>Number | Residue<br>Name |
| --- | --- |
| 353 | LEU |
| 351 | GLY |
| 399 | ARG |
| 396 | GLU |
| 414 | THR |
| 401 | GLU |
| 404 | SER |
| 398 | SER |
| 395 | GLU |
| 321 | THR |
| 324 | GLY |
| 352 | GLU |
| 350 | SER |
| 373 | GLU |
| 372 | HIS |
| 335 | GLY |
| 331 | GLU |
| 371 | SER |
| 332 | ASP |
| 368 | ARG |
| 391 | GLN |
| 403 | ASN |
| 343 | ALA |
| 407 | ASN |
| 405 | ARG |
| 397 | TYR |
| 333 | GLY |
| 369 | ASN |
| 380 | LYS |
| 376 | ALA |
| 381 | ASN |
| 377 | ILE |
| 409 | SER |
| 408 | SER |
| 354 | ARG |
| 392 | TYR |
| 363 | ASN |
| 382 | ALA |
| 318 | ARG |

**Table S2.**
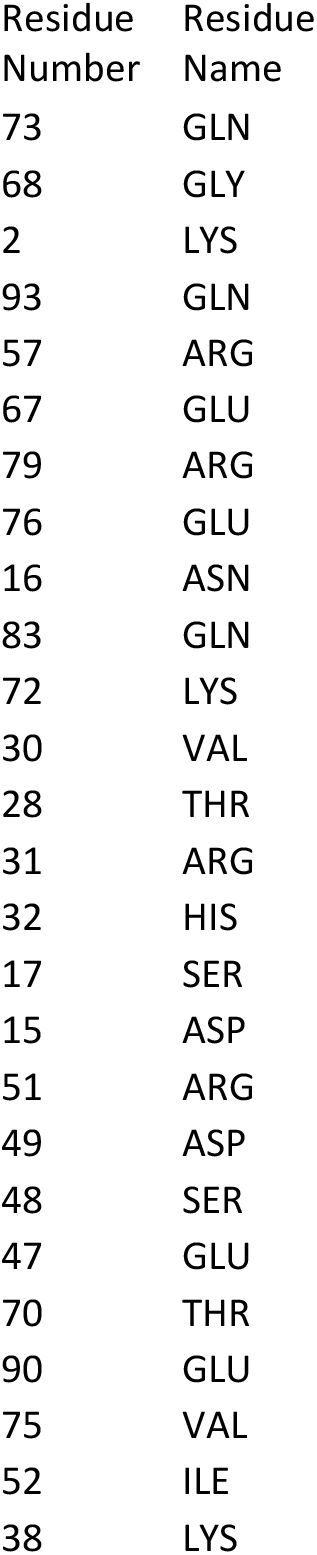
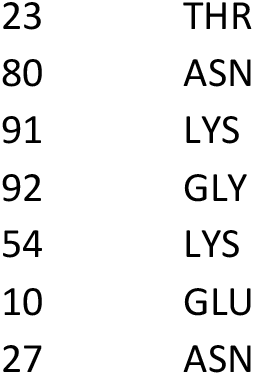
First 33 unique residues (35 percent of total number of protein residues) obtained from first principal component of hydrogen bonds for PDZ2 domain.

**Table S3.** First 55 unique residues (35 percent of total number of protein residues) obtained from first principal component of hydrogen bonds for Pin1 domain.

| Residue<br>Number | Residue<br>Name |
| --- | --- |
| 147 | SER |
| 159 | ILE |
| 115 | SER |
| 113 | CYS |
| 80 | ARG |
| 63 | LYS |
| 154 | SER |
| 131 | GLN |
| 104 | GLU |
| 102 | ASP |
| 21 | ARG |
| 34 | TRP |
| 157 | HIS |
| 151 | PHE |
| 109 | GLN |
| 107 | ALA |
| 59 | HIS |
| 38 | SER |
| 33 | GLN |
| 119 | ARG |
| 68 | ARG |
| 153 | ASP |
| 74 | ARG |
| 112 | ASP |
| 35 | GLU |
| 108 | SER |
| 36 | ARG |
| 37 | PRO |
| 111 | SER |
| 142 | ARG |
| 137 | ALA |
| 134 | PHE |
| 87 | GLU |
| 84 | GLU |
| 97 | LYS |
| 93 | ILE |
| 78 | ILE |
| 76 | GLU |
| 18 | SER |
| 23 | TYR |
| 56 | ARG |
| 121 | ASP |
| 77 | LYS |
| 16 | SER |
| 15 | MET |
| 72 | SER |
| 75 | GLN |
| 120 | GLY |
| 118 | ALA |
| 140 | ALA |
| 105 | SER |
| 82 | LYS |
| 66 | GLN |
| 162 | THR |
| 98 | SER |

**Note S1. Predicted allosteric signal transduction pathways for PDZ3 system**

Pathway 1 starts from the αC helix (experimentally verified allosteric site) and consists of βB-βC loop, αB helix to the binding site, which is in accordance with our previous observations.

### Important residues part of Pathway 1

αC helix (GLU 401-ARG 399-SER 398-TYR 397-GLU 396-GLU 395) - βB-βC loop region (GLY 335-GLU 331-ASP 332 –GLY 333) - αB helix (SER 371-HIS 372-GLU 373-ALA 376-ILE 377 –LYS 380 – ASN 381 –ALA 382)

Pathway 2 starts from the αC helix and consists of αA loop and βA-βB loop ending in canonical binding site.

### Important residues part of pathway 2

αC helix (GLU 401-ARG 399-SER 398-TYR 397-GLU 396-GLU 395) - αA helix and nearby loop (ARG 354-LEU 353 –GLU 352 –GLY 351 –SER 350-ALA 343) - βA-βB loop (THR 321-GLY 319-ARG 318)

**Note S2. Predicted allosteric signal transduction pathways for PDZ2 system**

Pathway I connects the αB helix to the distal αA helix (known allosteric site) via βA–βB loop

αA helix and nearby loop (ARG 51-ASP 49-SER 48-GLU 47) - βA–βB loop (SER 17-ASN 16-ASP 15) – αB helix residues(GLN 83-ASN 80-ARG 79-GLU 76-VAL 75-GLN 73-THR 70)

Pathway II connects distal N terminal region with αB helix residues

N terminal region (GLN 93-GLY 92-LYS 91-GLU 90) - βB–βC loop (ASN 27-THR 28-VAL 30-ARG 31-HIS 32)-αB helix residues(GLN 83-ASN 80-ARG 79-GLU 76-VAL 75-GLN 73-THR 70)

**Note S3. Predicted allosteric signal transduction pathways for Pin1 system**

### Pathway I

WW domain (MET 15 -SER 16 -SER 18 -ARG 21-TYR 23-TRP 34-GLU 35-ARG 36 – PRO 37 –SER 38) interdomain interface-PPIase domain core (α4 helix and β6-β7 loop region)(ARG 142-ALA 140-ALA 137

**-** LYS 132 - GLN 131 –THR 152-ASP 153-SER 154)–catalytic loop region (ARG 68-SER 72-ARG 74 – GLN 75-GLU 76-LYS 77-ILE 78)

### Pathway II

WW domain (MET 15 -SER 16 -SER 18 -ARG 21-TYR 23)-PPIase peripheral α1 helix, and the α1 helix-core interface (LYS 97-SER 98-ILE 93-ILE 89-PHE 151-THR 152-ASP 153)-to the catalytic loop (ARG 68-SER 72-ARG 74 – GLN 75-GLU 76-LYS 77-ILE 78).

